# Uncertainty drives deviations in normative foraging decision strategies

**DOI:** 10.1101/2021.04.24.441241

**Authors:** Zachary P Kilpatrick, Jacob D Davidson, Ahmed El Hady

## Abstract

Nearly all animals forage, as it is essential to acquire energy for survival through efficient search and resource harvesting. Patch exploitation is a canonical foraging behavior, but a systematic treatment of how animals cope with uncertainty is lacking. To address these shortcomings, we develop a normative theory of patch foraging decisions, proposing mechanisms by which foraging behaviors emerge in the face of uncertainty. Our model foragers statistically and sequentially infer patch resource yields using Bayesian updating based on their resource encounter history. A decision to leave a patch is triggered when the certainty of the patch type or the estimated yield of the patch fall below a threshold. The timescale over which uncertainty in resource availability persists strongly impacts behavioral variables like patch residence times and decision rules determining patch departures. When patch depletion is slow, as in habitat selection, departures are characterized by a reduction of uncertainty, suggesting the forager resides in a low-yielding patch. Uncertainty leads patch-exploiting foragers to overharvest (underharvest) patches with initially low (high) resource yields in comparison to predictions of the marginal value theorem. These results extend optimal foraging theory and motivate a variety of behavioral experiments investigating patch foraging behavior.

## 1 Introduction

Foraging is performed by many different species^1–5^ and engages cognitive computations such as learning of resource distributions across spatiotemporal scales, route planning, and decision-making^6^. Comparing species, one can ask how these integrated processes have been shaped by natural selection to optimize returns in the face of environmental and physiological constraints^6,7^. Foraging thus provides the opportunity to study and quantify how both evolution and neural circuitry shape a natural behavior^8–11^.

In natural landscapes, foraging involves a decision hierarchy that unfolds across multiple length and time scales, that consider both *where to forage* as well as *how long to exploit a certain resource*^12,13^. On long timescales, animals accumulate evidence to choose which of a collection of large areas they will dwell in and forage, during which their activity does not appreciably change the resource landscape. Following previous work, we refer to this as “habitat choice”^12^. Within habitats, animals exploit resources on shorter time scales, while their activity depletes resources in visits to localized regions. We refer to this behavior as “patch-exploitation”, or “patch-leaving”. Questions of where to forage and how long to exploit local patches of resource comprise a multi-level framework for examining behavior across spatial scales; still larger scales consider the home range of an individual, as well as the species range^13,14^. For both habitat-choice and patch-leaving, the local regions of an environment can be conceptualized as a “patch”; the key difference is in whether or not the forager’s activity impacts the resource availability in the landscape. As a decision problem, both are sequential choice processes, so the forager does not make a choice between discrete alternatives that are presented simultaneously, but rather only receives evidence from the current patch, and must decide whether to stay or go^15^. Thus, habitat-choice and patch-leaving are related but differ in the length and time scales involved (Fig. 1A). However, most theoretical work has considered these two problems separately.

**Figure 1.**
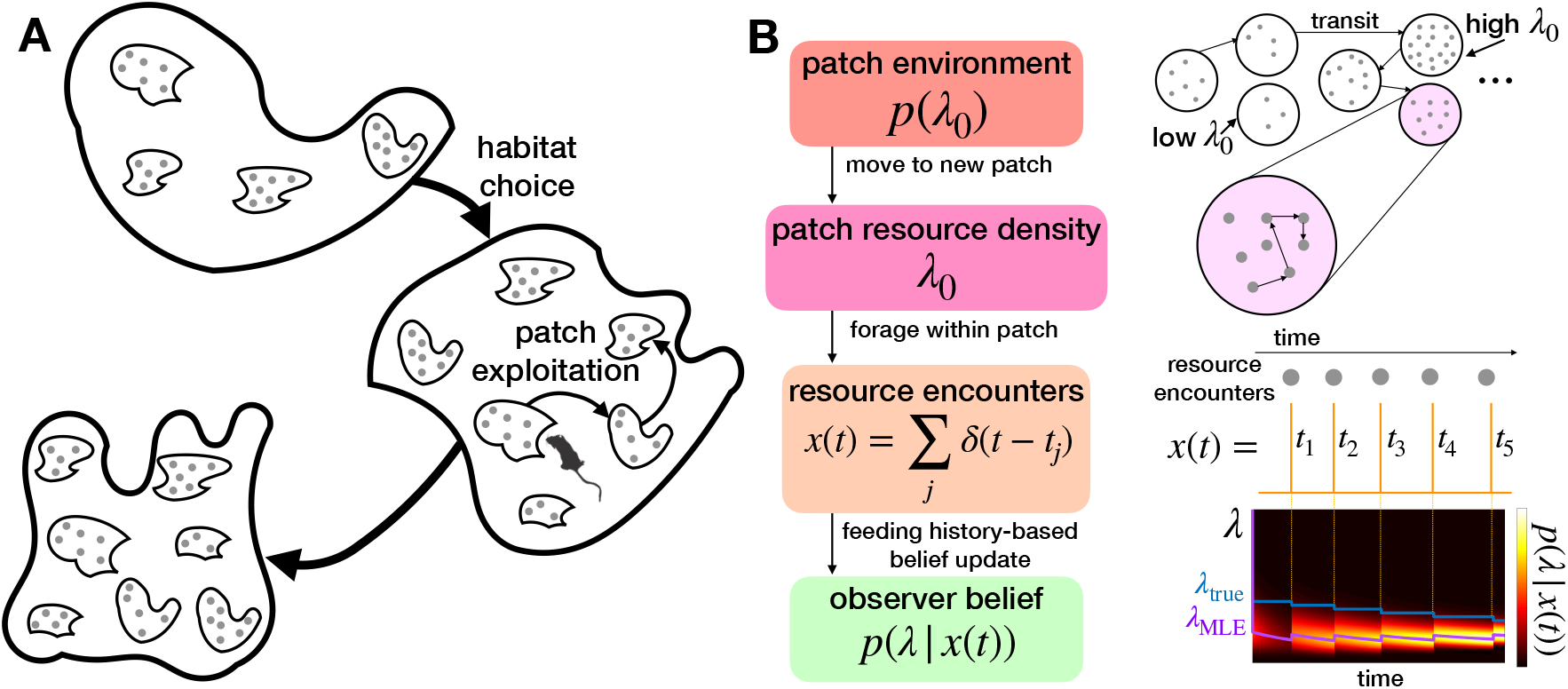
Patch-departure tasks and model. **A**. Task environments: On long timescales, an animal decides between habitats whose resource yields change slowly, and on shorter timescale, the animal exploits patches whose resources are depleted more quickly. **B**. Ideal observer foraging model: Initial yield of patch is drawn from the distribution *p*(*λ*_0_), generating random resource encounter times *t*_1:*K*_, and updating belief of current resource yield rate *λ* (*t*) for patch. The maximum likelihood estimate *λ*_MLE_ approaches the true *λ*_true_ yield rate over time.

Regarding habitat-choice, the optimal behavior is clear: the forager should locate and spend as much time as possible in the habitat patch with the best resources. Observational studies of habitat-choice often consider the combined effects of multiple factors to ask how well they predict observed habitat use (e.g.^16^). Theoretical and experimental studies have investigated multiple factors, such as density-dependent effects predicted by the ideal-free distribution when there are multiple foragers on a landscape^17^, and how perceptual constraints may lead to deviations from optimal choices^18^. However, with a focus on examining and predicting optimal behavior, most studies do not consider the process by which individuals use information to reach decisions of habitat-choice.

Patch-leaving considers shorter timescales, where the forager substantially depletes the patch during their visit, and must subsequently decide when to leave the current patch in search of another. A basic result in behavioral ecology, the marginal value theorem (MVT) states that an animal can optimize resource intake rate by leaving its current patch when the estimated within-patch resource yield rate falls below the global average resource intake rate of the environment^19^. While this theory has been validated in multiple behavioral studies^2–290^, it does not describe how animals learn key environmental features (e.g., resource distribution) from experience, or how such a decision rule can be implemented mechanistically. Most mechanistic models of foraging do not consider the process of statistical inference^20,21,25,28,30,^ and thus cannot explain how optimal foraging decisions are shaped by the presence and reduction of uncertainty based on resource encounters. On the other hand, Bayesian models of patch leaving that do ask how animals use limited information to make foraging decisions^31–33,33–48^ tend to consider a narrow range of environmental conditions.

The aim of our study is thus to develop a Bayesian framework of foraging behavior, treating decisions as a statistical inference problem, and connecting normative theory of foraging decisions with mechanistic evidence accumulation models^30^. We first define a general mathematical framework to model patch-foraging decisions that applies at different timescales with regards to search and depletion of the resource. These represent the different ecological decision cases described above: “habitat-choice” and “patch-leaving”. Because these cases are only separated in the timescale of resource depletion, we treat both with the framework of patch foraging as an evidence accumulation process whereby a threshold on available evidence triggers a decision.

Using several mathematically tractable cases in which probabilistic updating based on the receipt of resources within a patch can be modeled by stochastic differential equations (SDEs), we determine patch leaving statistics via solutions to first passage time problems. We thus obtain analytical expressions for optimal decision thresholds that connect to observable quantities of interest, including patch residence time, travel time, resource consumption over time, and patch yield rate over time. For both habitat-choice and patch-leaving, we show that in uncertain or resource-poor environments, uncertainty causes even an ideal Bayesian observer to tend to stay too long in low yielding patches (overharvesting) and not long enough in high yielding patches (underharvesting). Other studies have suggested that such a sub-optimality arises due to a limit in discriminating between similar patches^18^, or deviations from the MVT due to state-dependence or other decision factors^49^, but here we arrive at this result from principles of statistical inference. By establishing a general Bayesian framework for patch foraging at multiple scales, our study provides a platform to study behavioral and neural mechanisms of naturalistic decision-making akin to how trained decision-making behavior is studied within systems neuroscience^8,50^.

## 2 Sequential sampling model framework

The patch foraging model framework, which describes both habitat-choice and patch-leaving, considers an animal searching its environment, which contains distributed resource patches (Fig. 1A). When the animal enters a patch, they consume resources within the patch. We represent the decision process via a sequential sampling model for an ideal observer’s posterior of their current patch’s yield rate λ (t). This assumes an animal learns over time the yield of the patch they are currently in, in order to make a decision about if and when they should leave and search for another patch.

The initial yield rate 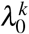 determines the rate at which the animal initially encounters a resource in the patch and is drawn from the distribution *p*(*λ*_0_). We assume the forager knows and initializes their belief with the prior *p*(*λ*_0_) when arriving in a patch (Fig. 1B). This simplifying assumption allows us to obtain tractable solutions. We later discuss future extensions that could explore how learning *p*(*λ*_0_) shapes long term foraging strategies. Considering randomly timed resource encounters within a patch, we use a Poisson-distributed rate *λ* (*t*) = *λ*_0_ − *ρK*(*t*) (as in random search^51^) that decreases with *K*(*t*), the number of resource encounters so far, where *ρ* is the impact of each resource encounter on the underlying yield rate of the patch. Resource encounter history can be described by the summed sequence of encounters, each at time 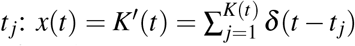. An ideal forager performs a Bayesian update of their belief about the current patch yield rate *λ* :

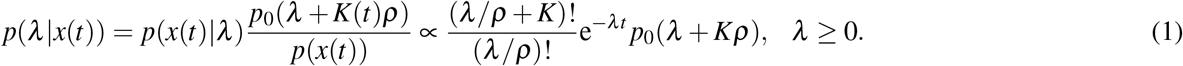

In general, resource encounters both: (1) give evidence of higher yield rates *λ*, since encounters are more probable in high yielding patches; and (2) deplete the patch, decrementing the yield rate *λ* by the *ρ* (Fig. 1B).

Changing *ρ* changes the rate of patch-depletion relative to the timescale of the foraging process. Small relative values of *ρ/λ*_0_ represent a large resource patch that the forager depletes very slowly. The limiting case *ρ/λ*_0_ → 0 represents the habitat-choice problem. Alternatively, when this ratio is intermediate up to unity (*ρ/λ*_0_ = 0.01 to 1), the forager considerably depletes the patch with each encounter. We refer to this as the patch-leaving problem, and show that in such cases, uncertainty in the patch yield can play a major role in shaping the departure strategy. We first consider the habitat-choice problem in Section 3, then following this, we consider the patch-leaving problem in Section 4.

## 3 “Habitat-choice”: Minimizing time to find high resource habitats

Habitat-choice refers to patch use at scales where the forager’s activity does not significantly affect the resource distribution or, in other words, that resource depletion occurs very slow relative to the time needed to the search process. We represent this with the theoretically tractable yet representative limit of zero patch depletion. In this case, the optimal behavior is to quickly locate a patch with the highest yield and remain there. Although in real environments, habitats eventually deplete and the forager would leave, our theoretical treatment of a “remain in the high patch” strategy can simply translate to a “stay a long time in the initially high yielding patch” strategy, with results applying similarly to both due to the separation of timescales: for habitat-choice, the time needed to search and decide on a high yield patch to remain in (which we denote as *T*_*arrive*_), is much less than the time that would be needed to actually deplete the patch.

Upon entering a patch, the forager must use their experience of resource encounters to decide whether to stay in the patch, or leave for another. We first consider a simplified *binary* environment where there are only two patch types – high yield versus low yield – and that the forager knows these possible patch types and their return rates. Here, the optimal behavior is to infer whether or not they are currently in a high yield patch, and if so, to stay, but otherwise to leave. Uncertainty and stochasticity of resource encounters means that the forager will visit some low yielding patches until they learn the yield rate and depart, and may also visit and depart from high yielding patches if the type is incorrectly inferred. We then consider more general cases, and show that the general trends and optimal strategies from the simpler binary case still apply; this includes environments with multiple patch types and environments with continuous distributions of patch types where the forager has a threshold for accepting a patch as sufficiently good. With this approach, we can explicitly derive statistics associated with patch departures and examine how the efficient identification of high quality habitats depends on environmental parameters like patch discriminability (e.g., *λ*_*H*_*/λ*_*L*_) and high patch prevalence (*p*_*H*_).

### 3.1 Two patch types

An environment with two possible patch types – high-yielding or low-yielding – is a theoretically tractable case that gives insight into optimal decision strategies and their resulting behavioral observables. Here, the distribution of patch types is *p*_0_(*λ*) = *p*_*H*_*δ* (*λ* − *λ*_*H*_) + *p*_*L*_*δ* (*λ* − *λ*_*L*_): *H* denotes the higher-yielding patch and *L* denotes the lower yielding patch. As stated, we assume the forager knows the values *λ*_*H*_ and *λ*_*L*_ and uses these as prior information to infer the type of the current patch. Using the limit of slow depletion (*ρ* → 0) to represent habitat choice, the animal determines which patch type they are currently in using the log-likelihood ratio (LLR) 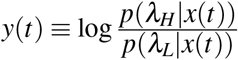. With this, their belief update can be written as a stochastic differential equation (SDE):

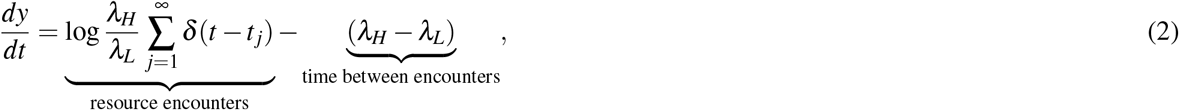

with initial condition set by the prior 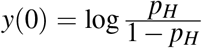. Elapsed time between resource encounters builds up evidence for the low-yielding patch while resource encounters provide evidence for the high-yielding patch. Eq. (2) has a simple form similar to classic evidence accumulation models of decision-making psychophysics^52,53^, recently extended to foraging decisions^30^.

The long-term resource intake rate is maximized if the forager finds and remains in a high-yielding patch (Fig. 2A). If the forager remains in a high-yielding patch, then the energy intake rate will reach *λ*_*H*_ in the limit of long time. Before locating and deciding to remain in a high-yielding patch, the forager may also visit low-yielding patches, leaving when their belief crosses the threshold (Fig. 2A), and may also visit and depart from high-yielding patches, if they are mistaken for low-yielding ones. The departure threshold sets the certainty that the forager obtains before leaving; a low threshold means high certainty of the patch type before leaving, while a high threshold will result in more departures. Too low of a threshold can lead to too much time spent in low-yielding patches while gathering more evidence, while too high of a threshold can lead to (incorrectly) departing from high-yielding patches before gathering enough evidence to distinguish their type. By setting the optimal threshold that balances uncertainty to minimize the time to arrive and remain in a high-yielding patch, we can ask how the environmental characteristics of relative patch yield, relative patch density, and travel time influence behavior and expected return of resources.

**Figure 2.**
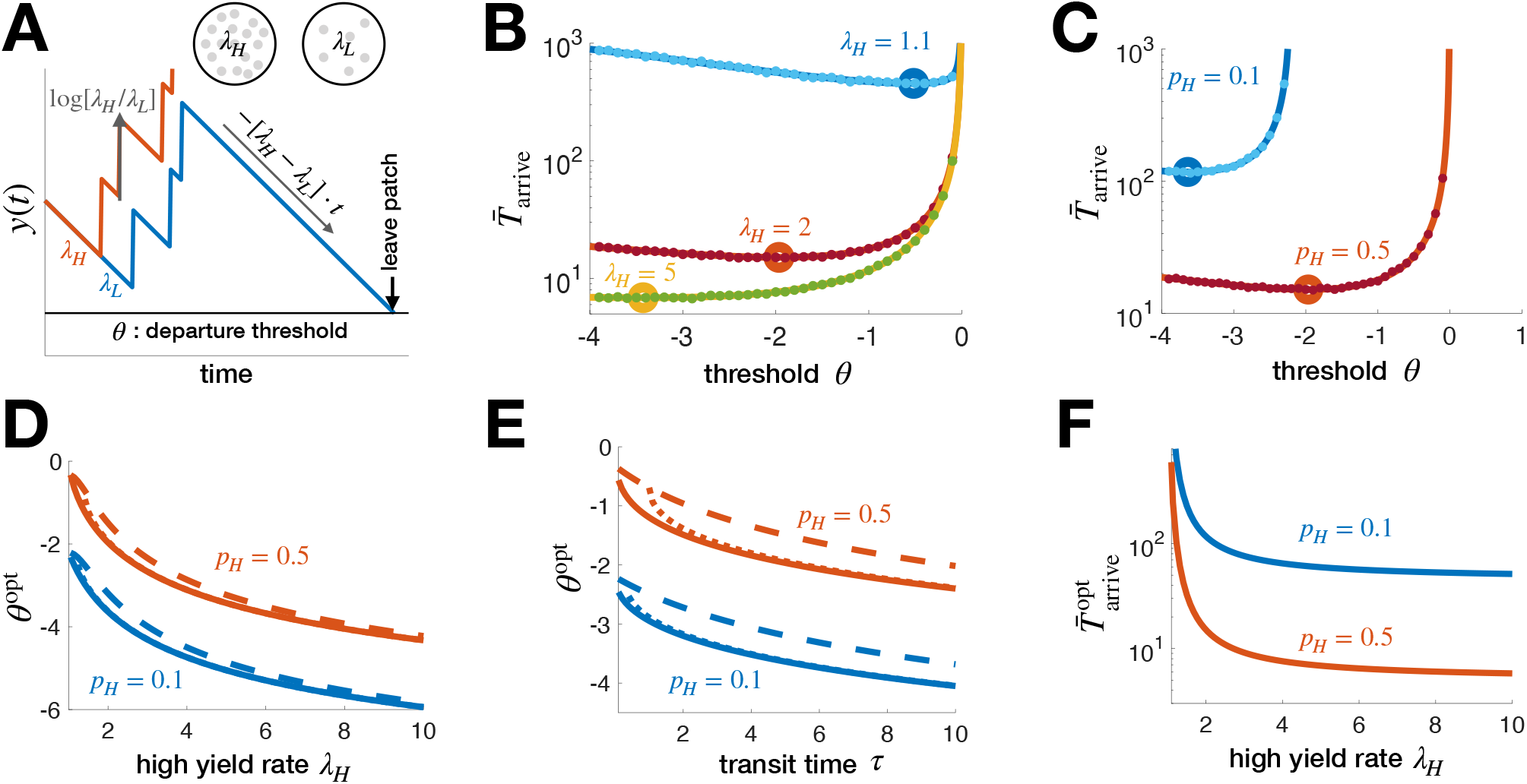
Statistics of habitat identification in environments with two patch types. **A**. Habitat type belief 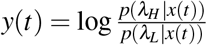 (as animal decides between a habitat with a high, *λ*_*H*_, and low, *λ*_*L*_, resource yield) increases with resource encounters and decreases between until *y*(*t*) = *θ* and the observer departs the patch. **B**,**C**. Mean time to arrive and remain in a high yield habitat varies nonmonotonically with departure threshold *θ* and decreases as the patch discriminability *λ*_*H*_*/λ*_*L*_ and high yield fraction *p*_*H*_ increase. Solid lines are Eq. (3). Dots are averages from 10^4^ Monte Carlo simulations. *λ*_*H*_ = 2 and *p*_*H*_ = 0.5 are fixed unless indicated. **D**,**E**. Departure threshold *θ*^opt^ minimizing the time to arrive in the high patch decreases with *λ*_*H*_*/λ*_*L*_ and *τ*. Solid lines are numerically obtained minima of Eq. (3), dotted lines are Eq. (4), and dashed lines are Eq. (5). In **D**, dotted lines appear overlaid on solid line because of the close fit. **F**. The minimal mean time 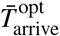 to arrive in a high yield patch decreases with *λ*_*H*_*/λ*_*L*_and *p*_*H*_. We fix *λ*_*L*_ = 1 and *τ* = 5.

The forager’s strategy is determined by the threshold *θ* on their belief (LLR), given by Eq. (2), shaping the time to find and remain in a high patch 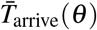. This quantity can be computed from the patch departure statistics of the forager, using first passage time methods, given the prior 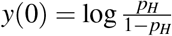 ^54,55^; *τ* is the mean travel time between patches which we assume is known or determined from experience. Using this, the time to arrive and remain in a high yield patch is

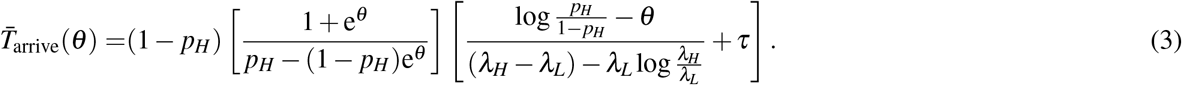

The minimum of this, corresponding to *θ*^opt^ (Fig. 2B,C) can be determined numerically (solid lines in Fig. 2D,E). An explicit approximation of *θ* ^opt^ is obtained by differentiating Eq. (3) dropping higher order terms and solving for

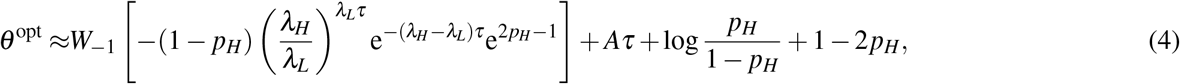

where 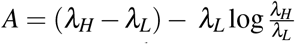 is defined for ease of notation (*A* = 0 for *λ*_*H*_ = *λ*_*L*_, and *A >* 0 for *λ*_*H*_ *> λ*_*L*_), and *W*_−1_(*z*) is the (−1)^th^ branch of the Lambert *W* function (inverse of *z* = *W* e^*W*^). This approximation matches well with the numerically obtained minima of Eq. (3) (dotted lines in Fig. 2D,E), and can be further simplified using the approximation *W*_−1_(*z*) ≈ log(−*z*) − log(−log(−*z*)) yielding

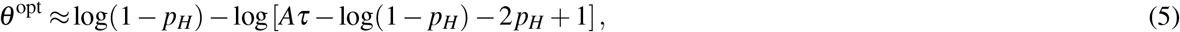

indicating the scalings of the optimal threshold in limits of environmental parameters (dashed lines in Fig. 2D,E). Details on the derivation of the optimal threshold can be found here^55^.

How best should an animal adapt their habitat search strategy to the statistics of the environment? When high yield patches are rare (low *p*_*H*_), travel times are large (high *τ*), or patches are easily discriminable (high *λ*_*H*_ relative to *λ*_*L*_) the forager should gain higher certainty by deliberating longer before departing a patch; indeed, from Eq. 5 and Figures 2D-E we see that the optimal threshold decreases with *ρ*_*H*_, *τ*, and *λ*_*H*_. Increasing discriminability (*λ*_*H*_*/λ*_*L*_) or the high yield patch fraction *p*_*H*_ decreases the minimal mean time 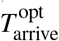 needed to arrive and remain in a high yield patch (Fig. 2F), since this makes finding a high yielding patch easier for the animal.

Furthermore, the outcome 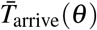 is most sensitive to the strategy (choice of threshold *θ*) in environments with low discriminability and a small fraction of desirable patches (Fig. 2B-C). In experiments, the value of *T*_arrive_ is an observable that can be used to infer the effective value of *θ* that an animal is using. The parameter sensitivity suggests an animal’s patch-selection strategy – i.e. the value of *θ* they are using – could be more precisely inferred when high patches are rare or more difficult to identify. Note, the patch-leaving rule of thresholding one’s LLR is mathematically equivalent to thresholding the mean estimated resource yield rate since 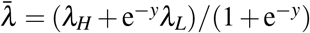, analogous to previous patch-departure rules developed^33,43^. Next, we generalize this approach to environments having more than two patch types, so decisions use multiple LLRs, such that optimal decisions do not simply map to thresholding estimated yield rate.

### 3.2 Multiple patch types

Animals may have to select from any number of patch types in an environment, which begs the question as to how decision and search strategies should extend to more general environments. With multiple patch types, decisions made by computing only two LLRs is sufficient to obtain near optimal performance in terms of minimizing the time to find and remain in a high-return patch. This result thus complements and extends previous work, which has considered optimal strategies for two patch types, or with fixed prior distributions^32,35,36^.

To model multiple patch types, consider environments with *N* patch types having resource yield rates *λ*_1_ *> λ*_2_ *>* … *> λ*_*N*_ ≥ 0 with patch fractions *p*_1_, *p*_2_, …, *p*_*N*_. Defining LLRs 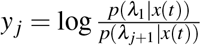 for *j* = 1, …, *N* − 1, yields the *N* − 1-dimensional system fully describing an ideal observer’s belief about the current patch type

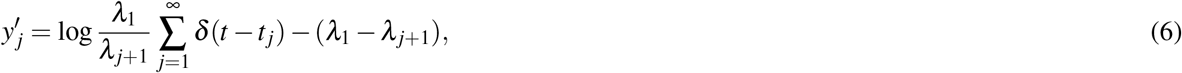

where 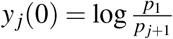, and any likelihood can be recovered as 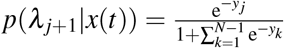, where *y*_0_ = 0 for *j* = 0.

As in the binary case, the optimal strategy is to find and remain in the highest yielding patch (*λ*_1_). We again represent patch-leaving decisions by thresholding the probability of being in the high yielding patch, such that when *p*(*λ*_1_ |*x*(*t*)) = *ϕ* ∈ (0, *p*_1_), the forager exits the patch. We approximate this thresholding process by requiring *y* _*j*_ ≥*θ* (for *j* = 1, 2, …, *N*) to remain in the patch, so the forager departs given sufficient evidence they are not in the highest yielding patch (See Fig. 3A for three patch types).

**Figure 3.**
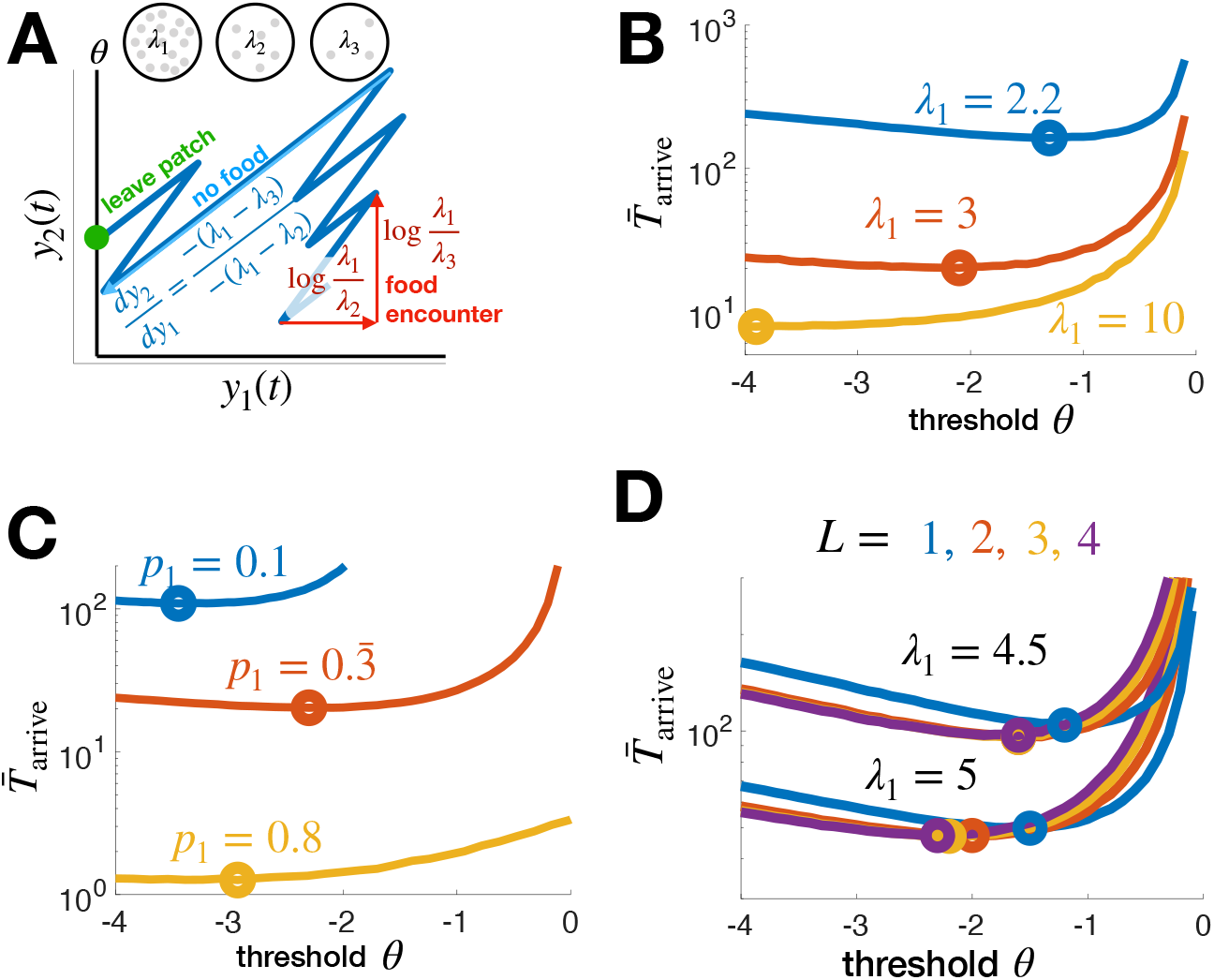
Optimal departure strategies for habitat choice in environments with multiple patch types. **A**. Beliefs about three possible habitat types 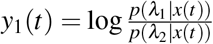 and 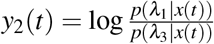 increase with resource encounters and decrease between until either reaches the departure threshold *θ*. **B**. Mean time 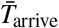 to arrive and remain in highest yielding patch *λ*_1_ (Eq. (7)) (with *N* = *L* = 3) decreases with patch discriminability *λ*_1_ as does the optimal departure threshold *θ* ^opt^ (circles). Three patch types with *λ*_2_ = 2, *λ*_3_ = 1, *τ* = 5, and *p*_1_ = *p*_2_ = *p*_3_ = 1*/*3. **C**. 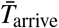 decreases with the prevalence of the best patch *p*_1_ (while *p*_2_ = *p*_3_ = (1 − *p*_1_)*/*2), but *θ* ^opt^ varies non-monotonically. **D**. The mean time 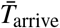 to arrive and remain in highest yielding patch *λ*_1_ (see Eq. (7)) depends on how many log-likelihood ratios (*L*) the forager uses to make a decision. Although the optimal time 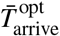 (curve minima – circles) decreases with *L*, the dependence is weak; *L* = 2 yields nearly identical mean optimal arrival times as *L* = 4. The optimal threshold which leads to 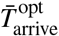 decreases with *L*. Other parameters are *p* _*j*_ = 1*/N, λ*_*j*_ = 6 − *j, j* = 1, 2, 3, 4, *N* = 5. In **C** and **F**, 10^6^ Monte Carlo simulations are used to compute the curves 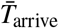.

To determine the sensitivity of strategies to their complexity, we also consider reduced strategies in which the observer only tracks the first *L* LLRs *y*_1_, *y*_2_, …, *y*_*L*_ and compares these with the threshold *θ* to decide when to leave the patch. Thus, we compute the mean time to arrive and remain in the high patch, which depends on the escape probability *π*_1_(*θ*) from the high yielding patch and the mean times to visit each patch 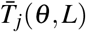 when escaping:

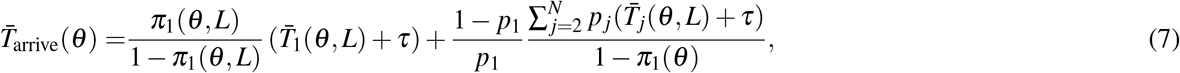

where patch departure strategy depends on the number *L* of LLRs thresholded and the threshold *θ* used.

The mean high patch arrival time 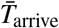 depends strongly on the high patch resource yield rate *λ*_1_, decreasing considerably as the patch becomes more discriminable (3 patches: Fig. 3B; 5 patches: Fig. 3D). On the other hand, 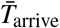 depends weakly on the worst patch’s yield rate *λ*_3_ (Fig. S1A), so uncertainty amongst the less valuable patches has little effect on behavior. In a related way, 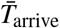 is much more strongly affected by changes in the fraction of the high yielding patch (*p*_1_: Fig. 3C) than changes in the balance of the middle (*λ*_2_) and low (*λ*_3_) patches (Fig. S1B).

Again, the optimal threshold decreases when patches are more discriminable: as *λ*_1_ increases the forager should gain a higher certainty before leaving (Fig. 3B). The average high patch arrival time 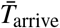 depends weakly on the threshold near the optimum, but the optimum threshold shows a non-monotonic dependence on *p*_1_. The lower optimum threshold for both low and high *p*_1_ values represents that in these cases it is optimal to be more certain of the patch type before leaving: for low *p*_1_ this occurs because high yield patches are rare (and thus there is a higher premium on distinguishing the high yielding patch when actually in one), and for high *p*_1_ this occurs because they are plentiful (one is more likely to land in a high yielding patch, so one can afford to require more certainty to depart). Between these cases, although the optimal threshold is slightly higher, the dependence is weak. Additionally, this demonstrates that if the forager did not know *p*_1_ (we assumed this is known and is used to formulate the leaving decisions), the best strategy would be to err on the side of choosing a low threshold, because the sensitivity of 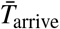 to threshold is relatively weak for choices too low, but can be higher for choices too high ((Fig. 3C).

In an environment with five patches, performance depends weakly on how many LLRs are used to make patch leaving decisions for *L >* 2. It is sufficient to simply track the LLRs between the first three patches, which correspond to using *L* = 2 (Fig. 3D). This is because the forager only needs to know whether they are in one of the best patches or not, since the goal is to eventually settle in one such patch as a habitat. This demonstrates again that the key features of uncertainty that matter to the optimal forager are the discriminability and prevalence of the best and second best patch type.

### 3.3 Continuum limit: many patch types

Building on the *N*-patch case, we now consider a scenario where there is a continuous distribution of patch qualities (*N* → ∞), so the resource yield rate for each patch *λ* is drawn from a continuous distribution *p*_0_(*λ*), which serves as a prior for the posterior *p*(*λ* | *x*(*t*)) with each patch visit (Fig. 4A). For any continuous probability distribution function *p*_0_(*λ*), the maximum *λ* will never be sampled, so arriving and remaining in the “maximum” yielding patch is not possible. We therefore assume the forager seeks patches with yield rates *λ*_*θ*_ or above, but deems lower yield rates to be insufficient. With this formulation, the forager updates a LLR based on a belief of whether current patch is greater or less than *λ*_*θ*_. Because this divides the continuous distribution into two categories, the mathematical treatment is similar to the binary case, but with added uncertainty because the patches in each category do not have the same return.

**Figure 4.**
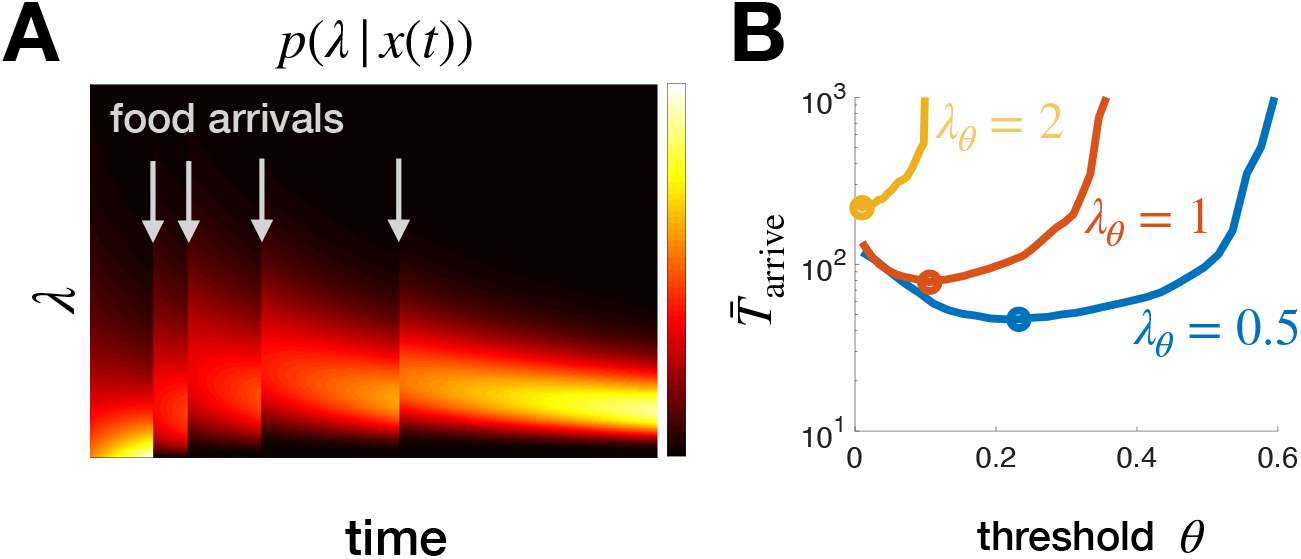
Habitat selection given a continuum of patch types. **A**. The posterior *p*(*λ* | *x*(*t*)) of the estimated yield rate is shifted up by each resource encounter and decreases between. **B**. There is an optimal *θ* that minimizes the time to arrive and remain in a high yielding patch (*λ > λ*_*θ*_), and the optimal time 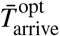 (circles) increases as the acceptable yield rate is increased. *α* = 1 is used here. 10^6^ Monte Carlo simulations for each *λ*_*θ*_and *θ*.

To model this, given a reference yield rate *λ*_*θ*_, we represent decisions in an environment with a continuous distribution of patch qualities by tracking 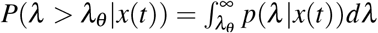. For the case of an exponential prior *p*_0_(*λ*) = *α*e^−*αλ*^, given *K*(*t*) resource encounters, we define 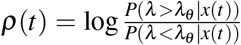 and state the forager departs the patch when 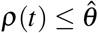 or when 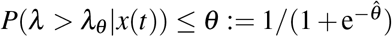. Note, to allow evidence accumulation, we require that 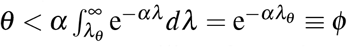, which represents the fraction of patches where *λ* ≥*λ* _*θ*_.

Computing the probability of escaping a high patch, 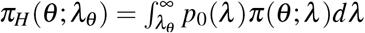, and the mean time per visit to a high and low patch types, 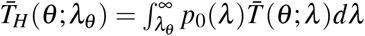 and 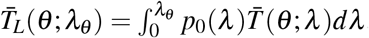, when departing (using Monte Carlo sampling), we then use Eq. (3) to compute the time to arrive in a high yielding patch (Fig. 4B). Placing a higher threshold *λ*_*θ*_ on the quality of an acceptably high yielding patch increases the time to arrive in an acceptably high patch. Moreover, the optimal threshold *θ* decreases, as more time must be spent in patches to discriminate a high yielding patch, which become rarer as *λ*_*θ*_ increases. Increasing *λ*_*θ*_ corresponds to making sufficiently high patches more discriminable and more rare.

With this formulation, the mathematical treatment in the case of a continuous distribution of patches is then the same as the binary case, and we can map corresponding results: setting a higher *λ*_*θ*_ is equivalent to decreasing *p*_*H*_ and concurrently increasing *λ*_*H*_. Although we considered an exponential distribution for *p*_0_(*λ*), we note that if this distribution changes, then this will affect the relationship between *λ*_*θ*_ and the equivalent mapping onto the binary case (in terms of *p*_*H*_ and *λ*_*H*_). Another further possibility would be to further “bin” the continuous distribution to correspond to 3 effective types, instead of two, as we did using a single threshold. The continuous case then would be treated analogously to a 3 patch type environment, and could be represented with two LLRs. However, further binning may not be necessary to achieve near-optimal decisions. Overall, this demonstrates that effective strategies for foraging environments with a continuum of patch types could be generated using particle filters that compute likelihoods over a finite set of patch types^56^.

### 3.4 Summary of results: Habitat choice problem

In general, we see that the results of considering the simple case of two distinct patch yield rates informs the general strategy when there are many possible patch types. The optimal time to arrive and remain in the highest yielding patch decreases as the high patch discriminability increases and as high patches become more common. Considering more than two patch types, the associated foraging strategies are most strongly coupled to environmental parameters of the highest and second highest yielding patch types. It is not necessary to compute LLRs associated with all possible types in order to efficiently find a high patch – even considering only a single LLR gives reasonable results, and the average time to arrive in a high patch is not strongly affected when the number of LLRs continues to increase beyond two. This suggests that animals select habitats by estimating a possible range of high-quality patches and then making patch-departure decisions based on whether patches meet those criteria or not.

## 4 Patch-leaving: Depletion- vs. uncertainty-driven decisions

When the scale of a patch is smaller, the forager will significantly deplete the patch’s resources during their visit. The decision is then not of which patch to remain in, but rather of when to leave the current patch in search of another. We therefore refer to this as patch-exploitation (Fig. 1A;^12,57^). While the nature of the resource differs for different animals (and typical patch residence times can accordingly vary from seconds up to hours - for some examples, see^31,38,47,58,59^). These cases all have in common that each resource patch is small enough such that availability within the patch is affected by the consumption of the forager.

The marginal value theorem (MVT) sets the optimal time to leave a patch in order to maximize resource consumption over time: when the current patch yield rate equals the overall average yield rate for the environment^19^. However, the MVT is simply an optimal rule, and does not specify the mechanistic process of how an animal uses its experience to reach a decision to leave a patch. Previous work has demonstrated that rewards in discrete chunks – instead of as a continuous rate – can affect the process an animal uses in decision-making ^21,24,30^. From a Bayesian perspective, decisions should use available information about the resource distribution in the environment. If resource availability within a patch is discrete or uncertain, even an ideal observer may not be able to accurately infer the actual rate of return, and thus would not be able to implement the leaving rule prescribed by the MVT. Experiments show that while the general trends predicted by the MVT hold in many cases, animals often deviate from an MVT-predicted strategy^49^. Moreover, in cases where patches contain very few items (e.g. 0 or 1 resource chunks), reward is not described by a rate function, and the MVT leaving rule does not apply.

Here we consider an animal that encounters resources in discrete chunks and infers the state of the environment and subsequently acts. This allows us to ask when the MVT rule is actually optimal versus when it does not apply, when deviations from the MVT occur due to uncertainty, and how a forager can incorporate prior knowledge about the resource distribution in the environment to reach a patch-leaving decision. We first treat the simple case of homogeneous patch types to establish the basic theoretical approach. Then, we consider an environment with two patch types to show how the inference procedure affects decisions in different environmental configurations, which we refer to as the “depletion-dominated” versus “uncertainty-dominated” regimes.

### 4.1 Homogeneous environments

To show how discreteness of resources affects decisions^21,24^, we first consider the simple case of a homogeneous environment with a single patch type. An ideal forager with prior knowledge of the initial yield rate *λ*_0_ can track time and resource encounters to determine the current yield rate *λ* (*t*), and then depart the patch when the inferred value of *λ* (*t*) falls below some threshold *λ*_*θ*_. Prior knowledge of the initial patch yield can be used in order to infer

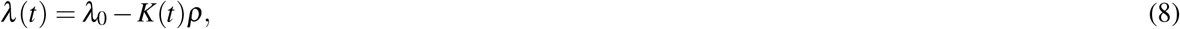

which represents the true underlying value of *λ* (*t*). Using this in a patch-leaving decision strategy is equivalent to departing after a fixed number of resource encounters^30^.

Using this inference strategy, we calculate the long term resource intake rate by assuming that *λ*_*θ*_ is an integer multiple of *ρ*. With this, the number of chunks consumed before departure is *m*_*θ*_ ≡ *K*(*T* (*λ*_*θ*_)) := (*λ*_0_ − *λ*_*θ*_)*/ρ*. Linearity of expectations allows us to compute the mean departure time as the sum of mean exponential waiting times between resource encounters 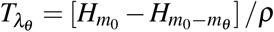, where *H*_*n*_ is the *n*th harmonic number. Thus, we can approximate the long term resource intake rate given *λ*_*θ*_ as

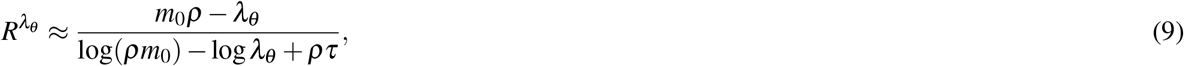

valid for *m*_0_ ≫ 1. There is an interior optimum *m*_*θ*_ that maximizes long term resource consumption rate, which we can estimate by computing the approximate critical point equation of Eq. (9)

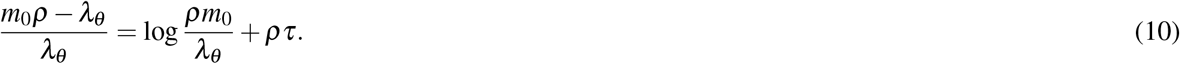

For large *m*_0_ (many chunks per patch), 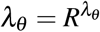, i.e. the leaving threshold is equal to the overall average rate of return in the environment; this is the optimum prescribed in the MVT^19^. Using the exact formula in Eq. (9), we can numerically determine the optimal threshold for small *m*_0_ (few chunks per patch). This shows that when there are only few chunks per patch, the true optimal threshold is close to, but not exactly equal to, 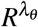 (Fig. 5A).

**Figure 5.**
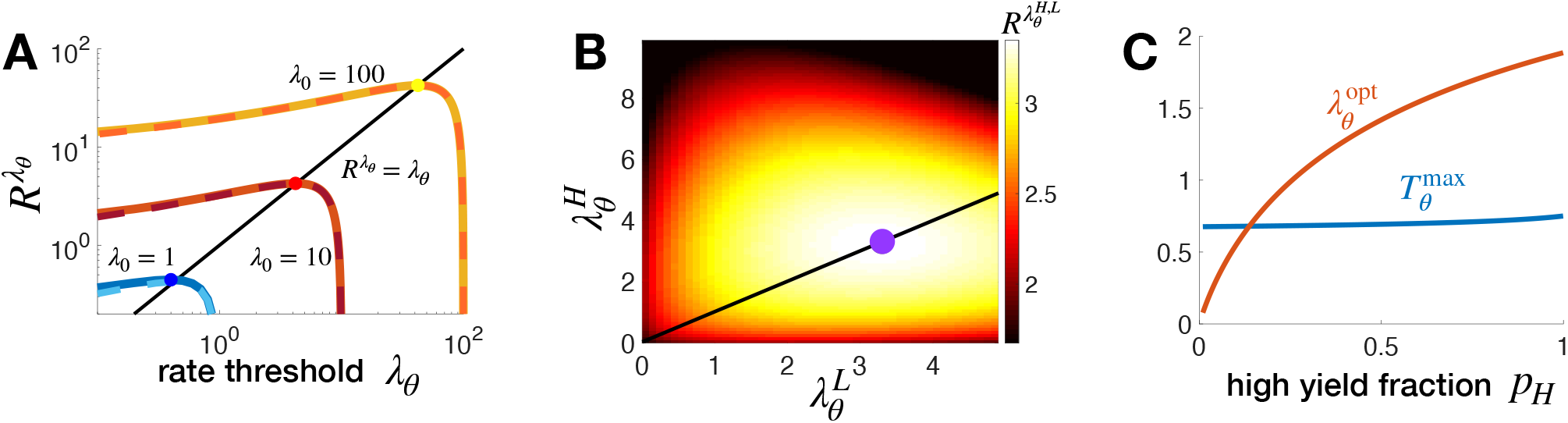
Departure strategies for the patch-exploitation problem. **A**. Rate of resource consumption as a function of strategy 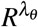, where the forager departs once the inferred yield rate falls to or below *λ*_*θ*_. Solid lines are exact solution of Eq. (9), and dashed lines are the large *m*_0_ approximation. Note optimal 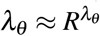. **B**. Rate of resource consumption 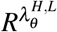 in binary environments in which the observer knows patch type (*λ*_*H*_ or *λ*_*L*_) upon arrival. Optimal strategy (purple dot) takes 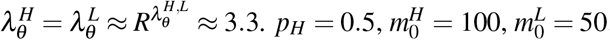, and *ρ* = 0.1. **C**. Optimal waiting time 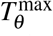 and departure threshold 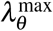 in a binary environment where the low patch has zero chunks of resource and the high patch initially has 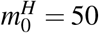. *ρ* = 0.1. Transit time *τ* = 5 throughout.

### 4.2 Binary environments

In binary environments, the forager estimates the underlying yield rate of the current patch, 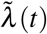, and departs when this falls below a threshold *λ*_*θ*_. Constant threshold strategies are our focus due to their relative simplicity, but we note that alternatively dynamic programming could be used to determine optima of a more general class of departure strategies^60^. We assume the forager uses knowledge that two different patch types exist to estimate the yield rate of the current patch; this involves using prior information to discriminate the patch type (high or low), combined with resource encounters which decrement the estimated yield rate. The belief can be determined according to a non-autonomous SDE for an LLR:

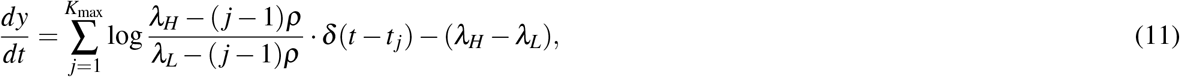

where 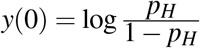 as in the case of habitat choice.

We focus on two different scenarios, which we refer to as: (a) Depletion-dominated regime, where the initial yield rate of the patch is known, and therefore leaving decisions are based solely on depletion, and (b) Uncertainty-dominated regime, where the type of the patch is not known upon entry, and optimal leaving decisions must consider uncertainty in the estimate of the current yield rate of the patch. For the uncertainty-dominated regime, we consider first the cases where low return patches contain zero resources, and then generalize to different amounts of resources per patch type.

#### Depletion-dominated regime

To represent what we term the depletion-dominated regime, we assume that the forager arrives in a patch and immediately knows the patch type *λ*_*j*_ (*j*∈ {*H, L*}) in which they reside (e.g., due to information provided by conspecifics or visual cues). In this case, the forager can make an accurate estimate of the true underlying yield rate of the patch, and therefore there is no uncertainty. Thus, leaving decisions are driven by depletion of the patch, as determined by when the estimated yield rate falls below some level 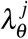.

Following our calculations from the homogeneous case, the long term resource intake rate depends on the initial resource chunk count in patches of type *j*, 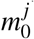, and the departure thresholds, so in the large 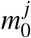 limit,

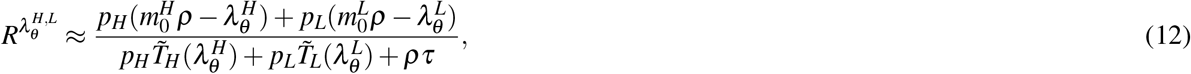

where 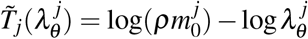 and the critical point equations for each partial derivative 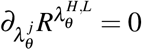 imply

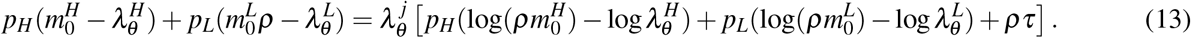

This can be rewritten as 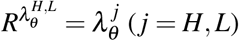. The aforementioned equation shows that, like the homogeneous case, an optimal strategy when there are many chunks per patch is to depart as the inferred yield rate equals the mean rate of resource encounters for the environment; additionally, the optimal threshold only depends on the average yield rate for the environment, and not the individual patch types (MVT: Fig. 5B). When there are few chunks per patch, the optimal threshold may slightly differ from this value (See results for the homogeneous case in Fig. 5A). The depletion-dominated regime is similar to a homogeneous environment: Since the forager knows the initial yield rate of the patch they are currently in, they can accurately infer the true underlying yield rate, and depart based on depletion of the patch. As in the homogeneous case, the optimal decision strategy can be formulated equivalently as either leaving when the estimated rate of return falls below a threshold, or as counting – leaving after consuming a certain amount of resources.

#### Uncertainty-dominated regime - Empty low patch

In the “uncertainty-dominated” regime, the forager does not know the initial yield rate of the patch upon entry. However, we assume that they have prior knowledge of the types of patches in the environment, i.e. that they know the values of *λ*_*H*_ and *λ*_*L*_. We first consider the tractable scenario where the low yielding patch is empty (*λ*_*L*_ = 0). Such situations occur if certain regions of the environment appear to have food (e.g., fruiting vegetation) but on closer inspection turn out to be empty (e.g., already foraged or rotten). The optimal strategy is for the observer to first wait a finite time *T*_*θ*_ to depart if no resources is encountered, but if resources are encountered before *t* = *T*_*θ*_, consume resources until the inferred yield rate drops to *λ*_*θ*_ = (*m*_0_ − *m*_*θ*_)*ρ*. Early/late decisions are thus driven by uncertainty/depletion. Assuming *m*_0_ ≫ 1, we can continuously approximate the long term resource intake rate and find it is maximized using a waiting time *T*_*θ*_ that is insensitive to *p*_*H*_. However, the threshold *λ*_*θ*_ depends on both *p*_*H*_ and travel time, because these parameters affect the overall average rate of resources available in the environment (Fig. 5C). The optimal threshold 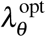 is not specified by the MVT, since uncertainty drives the forager to spend non-zero time in empty patches, adding extraneous time to the foraging process.

#### Many resource chunks

Next, we generalize to examine binary environments in which 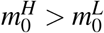 are arbitrary integers. In this case, the belief 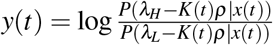 volves according to Eq. (11). The forager estimates the current yield rate of the patch from this belief,

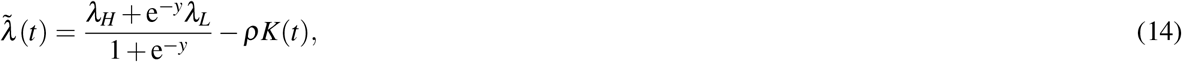

and an optimal strategy is to depart when 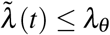. The threshold *λ*_*θ*_ should be tuned to 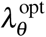 so the long term resource intake rate

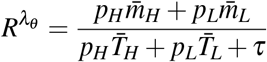

is maximized. We can compute departure times 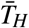 and 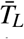 numerically via Monte Carlo sampling. For an environment where the overall availability of resources is low and there are few resource chunks per patch, the optimal strategy when the patch type is known is to fully deplete each patch before leaving - this is represented by a threshold of *λ*_*θ*_ = 0 for the inferred return rate. However, this is only optimal when the patch type is known; in the case of unknown patch type, the forager has uncertainty in whether or not there are remaining resources in the patch, and this causes the optimum threshold to be nonzero. In both cases, discreteness of resource encounters causes the optimal threshold to be lower than predicted by the MVT, although within this range of threshold values the average resource intake actually received is similar (Fig. 6A). In the case of high resource availability and many chunks per patch, the optimal thresholds are similar whether or not the forager knows the patch type upon arrival (Fig. 6B; compare black/blue curves), and coincides with optimal threshold predicted by the MVT.

**Figure 6.**
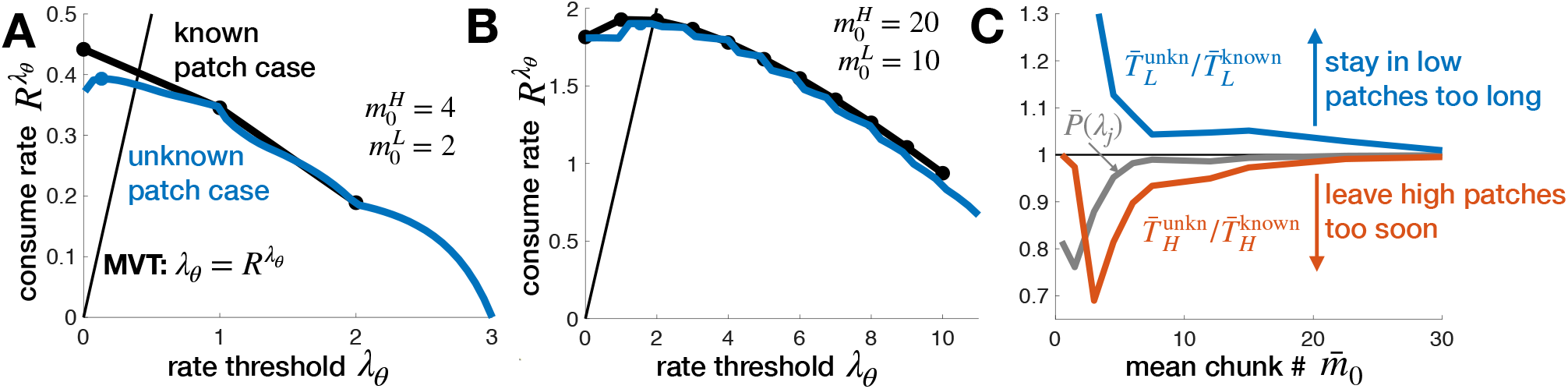
Depleting environments with ‘known’ vs. ‘unknown’ patch resource values. In the ‘known’ case, the forager knows the patch type and initial resource yield rate upon arrival, whereas in the ‘unknown’ case the forager must infer the path type. Patch leaving strategies in the unknown case approach those of the known case in the high chunk count limit. **A**. Total resource intake rate 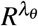 as a function of the estimated yield rate *λ*_*θ*_ at which the observer departs a patch. For low initial resource levels 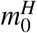 and 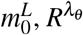 near the optimum (dots) deviates between the case of initially known patch type (black) vs. initially unknown (blue). Both thresholds are less than the MVT prediction due to discreteness of resource encounters. **B**. Resource intake rates of the known/unknown cases converge at higher initial resource levels. **C**. Deviations in optimal resource intake rate between known and unknown cases are due to less certainty 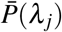 (grey line) about patch yield rate upon departure, leading the observer to stay in low patches too long (blue line) and leave high patches too soon (red line). Other parameters are *ρ* = 1, *τ* = 5, and *p*_*H*_ = 0.5. Mean rates and departure times in the unknown patch case were computed from 10^6^ Monte Carlo simulations.

Comparing cases, we see that foragers in sparser environments (lower average initial resource amount 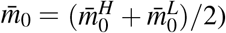 stay in low patches too long and leave high patches too soon in comparison to observers that immediately know their patch type, due to their uncertainty about their current patch before and at the time of departure (Fig. 6C). Uncertainty thus drives animals to underexploit (overexploit) high (low) yielding patches when high and low patches are different enough so that it is optimal to spend significantly more time in high versus low patches, but similar enough as to not be immediately distinguishable. We also note that in contrast to the depletion-dominated regime, where an optimal strategy can be implemented equivalently as either through rate-estimation or by leaving after consuming a certain amount of resources (counting). Optimal leaving decisions in the uncertainty-dominated regime must use a rate-estimation process, because of the associated uncertainty in the true yield rate of the current patch.

## 5 Discussion

Patch foraging is a rich and flexible behavior where an animal enters a patch of resources, harvests them, and then leaves to search for another patch. An animal’s behavior can be quantified by their patch residence time distribution, travel time distribution, the amount of resources consumed, and the movement pattern between patches. In this work we used principles of probabilistic inference to establish a normative theory of patch leaving decisions. With this general framework, we showed how foraging at different temporal and spatial scales are connected by a similar decision problem: “habitat-choice” refers to larger scales when foragers do not significantly deplete a resource, and “patch-exploitation” refers to smaller scales where the forager’s activity depletes the patch. For habitat-choice, the optimal behavior is to quickly locate and remain in a high-yielding habitat, while for patch-exploitation, it is optimal to use prior information along with reward encounters to estimate the current underlying yield rate to determine when to leave the patch (Fig. 7; Table 1). In ecological contexts, these activities are part of a behavioral hierarchy, where an animal must decide *where to forage* and *how long to exploit a certain resource*.

**Figure 7.**
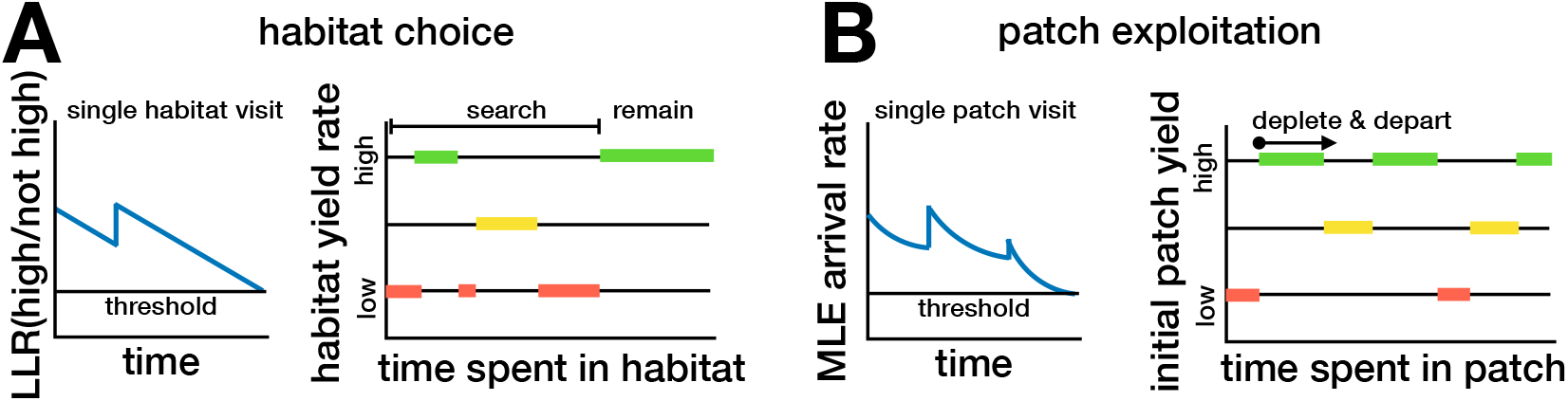
Summarized taxonomy of foraging strategies. See Table 1 for details. In different environments with three patch types (low: red, medium: yellow, high: green yielding), the different time series of decision variables (for a single patch decision) and patch visit time intervals. **A**. In habitat choice, an animal must determine whether their current patch is of the highest yielding type, departing if the probability they are not in the highest reaches some threshold, and undergoing a sequence of patch visits until finding and remaining in a high yielding patch. **B**. An ideal forager performing patch exploitation infers the yield rate of their current patch and departs when resource yield rate reaches a threshold, continuing patch visits indefinitely in large environments.

**Table 1.** Detailed taxonomy of departure decision strategies. Departure strategies and observable trends depend on environment and task: habitat selection or patch-exploitation (see also Fig. 7). Columns describe important aspects of the optimal decision strategy for each case, along with key model results.

| *Environment* | *Decision strategy and dependencies* | *Equations* | *Figures* |
| --- | --- | --- | --- |
| **Habitat selection.** |  |  |  |
| Objective: Minimize time to find highest yielding habitat. |  |  |  |
| Known: Resource yield rates of each habitat type. |  |  |  |
| $N$-habitat types | • Depart habitat when likelihood of being in highest yielding habitat falls below a threshold.<br>• Optimal strategy and arrival time depend on fraction and discriminability of high yield habitats. | $N = 2$: Eq. (2);<br>$N \geq 3$: Eq. (6). | $N = 2$: Fig. 2;<br>$N \geq 3$: Fig. 3 |
| Continuum of habitat types | • Categorize habitats as high or low yielding and depart habitat if likelihood of a high yield falls below a threshold.<br>• Time to identify high-yielding habitat is non-monotonic in departure threshold, and much longer when high patches are rare. |  | Fig. 4 |
| **Patch-exploitation.** |  |  |  |
| Objective: Maximize mean resource intake rate $R$ over a long time (several patches). | | | |
| Known: Initial yield rates of each patch type. |  |  |  |
| 1-patch type | • Depart when yield rate $\lambda(t)$ falls to a threshold value $\lambda_\theta$.<br>• Matches MVT except when there are very few chunks per patch in which case the forager should empty the patch. | Eq. (10) | Fig. 5A |
| 2-patch types: patch type known on arrival | • Depart when resource yield rate $\lambda_j(t)$ reaches a threshold.<br>• Represents ‘Depletion-dominated’ regime. Recovers MVT. | Eqs. (12) & (13) | Fig. 5B |
| 2-patch types: empty low yield patch | • Wait a time $T_\theta$, then depart patch if no resources found. If resources are encountered by $t < T_\theta$, use threshold on inferred yield rate to make leaving decision (similar to single patch type case).<br>• ‘Uncertainty-dominated’ regime deviates from MVT.<br>• Optimal wait time and departure threshold $\lambda_\theta$ increase with prevalence of high-yielding patch. |  | Fig. 5C |
| 2-patch types: both high and low patches have resources | • Decision via threshold on current estimated yield rate $\tilde{\lambda}(t)$. Choose optimal threshold $\lambda_\theta$ that maximizes long term resource intake rate.<br>• Optimal return differs from known case given few resources per patch, converges to known patch case when resource density is high.<br>• Forager stays in low patches too long, leaves high patches too soon when there are few resources per patch (uncertainty-dominated regime). | Eq. (14) | Fig. 6 |

In the case of habitat choice, the forager should use their experience of reward encounters to determine whether to stay or leave; in our model an optimal forager departs a habitat when their log-likelihood ratio for the probability of high yield versus other habitats falls below a threshold. Optimal decisions are based on inference of habitat quality, with uncertainty being the driving factor in habitat departure times; while this is related to resource intake rates, it is not the same, because of how prior information can be used in patch inference. We showed that with multiple different patch types, it is not necessary to track LLRs for all patch types – behavior is most strongly affected by inference related to the highest and second-highest yield habitat types. The optimal time to arrive and remain in a high yielding habitat is lower when patches are more discriminable, or when high yield patches are more common. While *T*_arrive_ is an experimentally observable quantity, an animal’s internal decision threshold is not; our model connects these quantities, and thus can be used to infer the decision rules an animal is using (Fig. 2). We showed that it is optimal to have a lower threshold – and thus gain a higher certainty before leaving – when travel times are large, high patches are rare, or high patches are easier to discriminate. These results give quantitative predictions that can be used to interpret experiments, for example, to examine whether the animal is minimizing the time to reach the highest yield habitat in a heterogeneous environment. Moreover, we showed how behavior in the general case where many habitat types exist can be understood by mapping results onto the tractable case of only two different patch types (Figs. 3 & 4). By varying systematically the percentage of high yielding habitats and the discriminability (ratio of high to low yield rate), the model predicts how this affects the minimal time to arrive at the highest yield habitat, and connects this to a process that could implement such computations.

For patch-exploitation, when the forager depletes patches in their habitat, in most cases the long term intake rate is maximized by departing a patch when the in-patch estimated resource yield rate matches the average return rate of the environment (i.e. by implementing the MVT rule). However, this does not apply when resources within a patch are limited so there is more uncertainty about the yield of the current patch upon departure (Fig. 6). Often in nature, the environment is volatile and animals make foraging decisions while uncertain about local resource availability^61^. Our model predicts that if there is high uncertainty about the patch type, this causes even an ideal Bayesian forager to stay longer in low yield patches and shorter in high yield patches than predicted by the MVT.

Our theoretical treatment of patch leaving decisions builds on previous Bayesian models of foraging^33–41,43.^ Our approach goes further than previous work by providing a step-by-step derivation of the normative strategies associated with a continuum of different environmental conditions, systematically identifying the dependence of observable behaviors (e.g., patch departure times) on environmental parameters. During habitat choice, the minimal arrival time to the high yield habitat scales with the probability of high yield patches in the environment. On the other hand, we have shown that in the case of depleting patches, the amount of time a forager overstays or understays in a patch scales with the density of resources in the patch. We are also able to infer the optimal threshold an animal should use. While this cannot be measured directly in experiments, our observations do reveal environmental parameter regimes under which performance (e.g., foraging yield) is sensitive to changes in strategy. This can not only inform the design of behavioral foraging experiments, so as to determine task parameters that best reveal an animal’s strategy, optimal yields can also be compared with those obtained by animals in the wild to see how finely tuned their foraging strategies are.

Analysis of experiments shows animals forage in ways that suggest they use Bayesian reasoning^42–48^, using prior knowledge of their environment to modulate foraging behavior^59,62,63^. For example, bumblebees^31^ and Inca doves^64^ adjust their foraging strategies in response to the predictability of the environment, as a Bayesian forager would, but this is not a universal trend^65^. Patch-leaving decisions may deviate from Bayes optimality as animals become risk-averse in variable environments^66^. Although other Bayesian models have considered patch-foraging decisions^32,36^, it has been difficult to relate these results to experiments due to limited theoretical considerations of the environmental configuration. By showing how trends from more realistic environmental distributions relate to the theoretically simpler case of two patch types, and suggesting a mechanistic implementation that the forager can use to implement optimal decision rules, our model provides optimal behavior predictions that can be used to interpret experimental results. Moreover, our theoretical approach applies not only to patch-exploitation, but also to habitat-choice – where the MVT does not apply – and thus enables connections across these multiples scales of behavior^13^.

Although we used a constant threshold value based on either the belief or estimated yield rate, other work has examined cases where optimal decision strategies involve time-dependent decision thresholds^60,67–69^. Typically, these results arise in the context of multi-trial experiments in which the quality of evidence on each trial varies stochastically and is initially unknown. In the habitat choice context, the quality of evidence is fixed across habitat visits, fitting the assumptions of classic, constant threshold optimal policies. We would therefore expect an analysis allowing for a dynamic threshold to yield the same results as we obtained here. On the other hand, when the animal performed patch-exploitation in an uncertain binary environments, we projected a higher-dimensional description of the patch value to a single scalar estimate of the patch yield rate. In this case, a constant-threshold implementation may not be purely optimal. Leveraging methods from dynamic programming commonly used to set optimal decision policies^68^ would be a fruitful next step in ensuring the optimality of our patch leaving decision strategies. A common theme in previous Bayesian models that use dynamic programming^32,43^ and our approach, is that the forager should use the expected future return, not the current return, to make departure decisions. An advantage to the constant-threshold treatment is that it enables simple explicit quantitative relations that can be used to interpret experimental data (Fig. 2). For example, experiments evaluating how animals value the quality of evidence could help us delineate whether the animal is using a constant threshold or not.

Effective search is integral to survival in nature^70^, and search behavior can give information on individual decision strategies. One can consider search within and in-between patches. Although we assumed random timings of resource encounters, an extension of our model could take into account different spatial arrangements. Within patches, an animal may perform random or a systematic search. For example, a recent study found that rats can solve the stochastic traveling salesman problem using a nearest neighbour algorithm^71^. A similar approach could ask how an animal’s search and navigation pattern interact with different patch leaving decisions to create an effective foraging strategy that also considers memory of specific patch locations. Indeed, the explicit consideration of spatial movement may be necessary to understand foraging decisions. Previous work found that when rats must physically move to perform foraging, the observed behavior differed from tasks that “simulate” foraging by presenting sequential choices or that consider visual search^72^. It is an open question as to how the animal integrates aspects of spatial movement with economic valuations of future reward.

Our model assumes animals know the initial yield rate of each type of patch in the environment. In a real life context or in an experimental setup, the animal would learn the environmental parameters, which we could model by considering another level in the inference hierarchy whereby the patch type quality and fraction are learned along with the transit time distribution. Although we considered a single forager acting alone, another important extension will be to consider interactions between animals, either through predator-prey interactions which affect foraging decisions^73^, social foraging of groups ^74,75^, or even competitive foraging^17,18,76^. In our model, the forager only receives direct (non-social) information about resource availability; in collective foraging, an individual receives both social and non-social information^77,78^. This can significantly affect foraging decisions, for example in the case where an individual must balance resource-seeking with group cohesion. Building on our modeling approach, foragers could share social information either by cooperating in the inference of the patch quality or by signaling to each other when to depart a patch as a threshold is reached.

To conclude, our model establishes a formal framework for the quantitative analysis of a natural behavior – patch foraging (involving both habitat-choice and patch exploitation) – that can be studied in the same formal rigor as many trained behavioral tasks. Such validated *behavioral algorithms* are crucial for the systematic design of future experiments and interpretation of data on animal behavior^79^. By comparing with theoretical optimal strategies, experiments and data can be used to understand the decision strategies an animal is employing and relate these to recorded animal movement and neural data. Future work will build on this model framework to generate testable hypotheses on the role of social interactions and the neural mechanistic underpinnings of foraging behavior.

## Acknowledgements

We thank Jan Drugowitsch for helpful discussions regarding the optimality of decision threshold policies across patch visits. ZPK was supported by a CRCNS grant (R01MH115557) and NSF (DMS-1853630). AEH acknowledges support by NIH grant 1R21MH121889-01. JDD acknowledges support by the Deutsche Forschungsgemeinschaft (DFG, German Research Foundation) under Germany’s Excellence Strategy - EXC 2117 - 422037984, as well as support from the Heidelberg Academy of Science.

## Appendix

**Figure S1.**
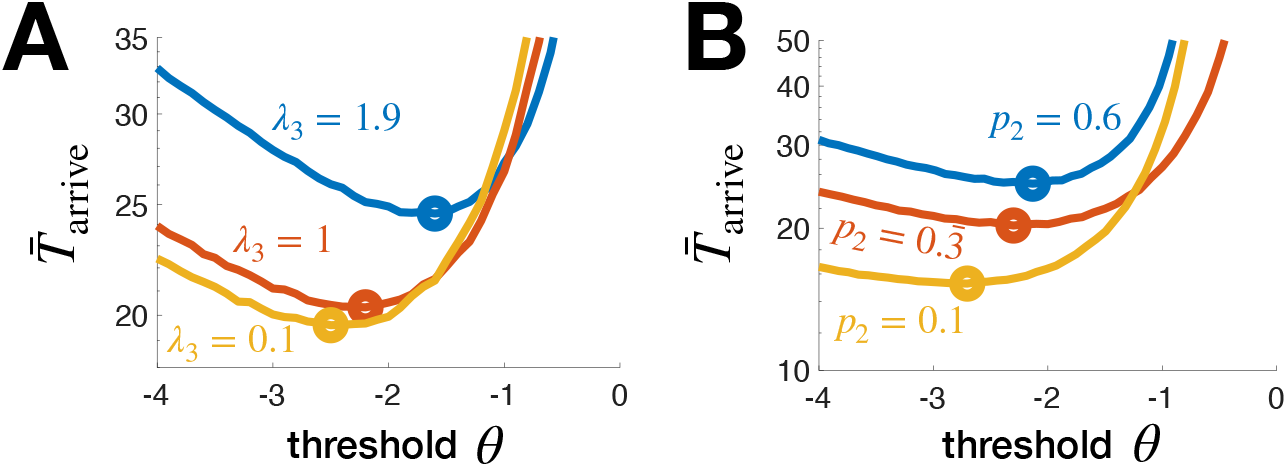
Insensitivity of patch finding performance to worst patch statistics. **A**. 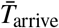 increases with *λ*_3_ as does *θ* ^opt^, since the worst patches become less easy to distinguish from the best (*λ*_1_ = 3). Here *λ*_2_ = 2, *τ* = 5, *p*_1_ = *p*_2_ = *p*_3_ = 1*/*3. **B**. 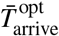 increases with *p*_2_ (while *p*_1_ = 0.333 and *p*_3_ = 1 − *p*_1_ − *p*_2_) as does *θ* ^opt^. All curves for 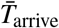 computed from 10^6^ Monte Carlo simulations.

